# Pandoomain, a scalable pipeline for genomic and protein domain context analysis, reveals widespread PT-TG domain architectural diversity and novel polymorphic toxins

**DOI:** 10.1101/2025.10.01.679605

**Authors:** Emanuel Becerra Soto, Adam J. Oliver, Marcos H. de Moraes

**Affiliations:** Department of BioSciences, Rice University, Houston, TX, USA; Biochemistry and Cell Biology Graduate Program, Rice University, Houston, TX, USA

## Abstract

The rapid expansion of bacterial genome databases presents significant opportunities for functional discovery, yet a large fraction of genes and protein domains remain uncharacterized. Analyzing genomic context and domain architecture are powerful approaches for functional inference, but existing tools often lack the scalability and integrated workflow required for high-throughput analysis. To address this, we developed Pandoomain, a Snakemake pipeline that automates the acquisition of genomes from NCBI, identifies proteins of interest using Hidden Markov Models (HMMs), and performs systematic domain annotation and gene neighborhood analysis. We demonstrate the utility of Pandoomain through a comprehensive analysis of the poorly characterized pre-toxin TG (PT-TG) domain across 347,289 bacterial genomes. Our analysis revealed 10,226 PT-TG-containing proteins organized into 312 unique domain architectures, highlighting their association with diverse interbacterial antagonistic systems, including the Type VI, Type VII, and CDI systems. By leveraging genomic context, we identified a novel variant of the WXG trafficking domain, termed W10XG, and subsequently discovered 24 new families of associated toxin domains. We experimentally validated six of these toxins, confirming that five are neutralized by their cognate immunity proteins. Furthermore, our analysis revealed a significant enrichment of mobile genetic elements near W10XG and WXG domains compared to other trafficking domains, suggesting these loci are hotspots for genomic diversification. Pandoomain is an accessible tool that enables systematic, large-scale exploration of protein domains, and our analysis of the PT-TG domain provides a rich resource for future investigations into the mechanisms and evolution of bacterial antagonism.

## Introduction

The continued expansion of public repositories, which now host millions of bacterial genomes originated from pure cultures and microbiomes, presents both a significant opportunity for discovery and a substantial analytical challenge. This expansion has not been followed by elucidation of biological functions, as 30% to 40% of bacterial genes and many protein domains remain uncharacterized[1–3]. To address this challenge, one powerful approach is the analysis of genomic context. This method relies on the principle that genes with related functions are often physically clustered on the chromosome in conserved gene neighborhoods or operons[4]. This genomic co-localization is a strong evolutionary signature of a functional linkage and has been instrumental in revealing the roles of novel proteins within complex biological processes, including metabolic pathways and protein secretion systems. A complementary approach is the analysis of protein domain architecture, the specific composition and organization of functional domains within a protein, which provides direct clues to its function.

Investigating genes and protein domains co-localization has been extremely successful in the discovery of novel functions of antimicrobial bacterial toxin domains and their biological functions. Previous foundational work identified dozens of novel toxins and trafficking domains predicted to be involved in interbacterial antagonism[5, 6]. Subsequent studies of new bacterial toxins through genomic analysis continues to yield valuable insights and offers comprehensive databases for the scientific community to explore in functional analysis[7–10]. Although the code used to generate the output of these studies is available in certain cases, it is often not structured to expand their scope across different genomes or protein domains, precluding their expansion and scalability.

A variety of bioinformatic tools exist to facilitate the analysis of protein domain architecture and genomic context. While the examples discussed here are representative of a larger landscape of available options, many of which are highly specialized for niche applications, they can be broadly categorized as either web-based servers or command-line pipelines. Web-based platforms such as KEGG[11], BioCyc[12], Interpro[13] web interface, the EFI-Genome Neighborhood Tool[14], and BRENDA[15] offer user-friendly graphical interfaces ideal for small-scale visualization but lack the scalability and customizability required for analyzing thousands of genomes. In contrast, command-line pipelines like Bactopia[16], ProkFunFind[17], and Domainator[18] provide the power and flexibility for high-throughput analysis, though they typically require significant bioinformatics expertise for setup and may not be able to be executed locally for vast datasets. A significant gap, therefore, exists for a tool that combines the scalability of command-line approaches with an accessible, automated workflow, as many existing solutions lack a cohesive, end-to-end framework that integrates data acquisition, taxonomic information, and systematic architecture comparison into a single, reproducible pipeline.

To address these limitations, we developed Pandoomain, an automated and reproducible pipeline focused on analyzing domain architecture and genomic context across large numbers of genomes extracted from the National Center for Biotechnology Information (NCBI). In this work, we describe the Pandoomain pipeline and validate its utility through a comprehensive analysis of the PT-TG domain. We chose this domain due to its poor characterization and its widespread distribution in polymorphic toxin systems[19, 20] (Fig. 1). This analysis serves not only to demonstrate the power of our tool but also to uncover novel features of the PT-TG domain family and its role in interbacterial antagonism. The development of Pandoomain and the wealth of data on polymorphic toxins are open to the research community and can power future investigation enterprises.

**Fig 1.**
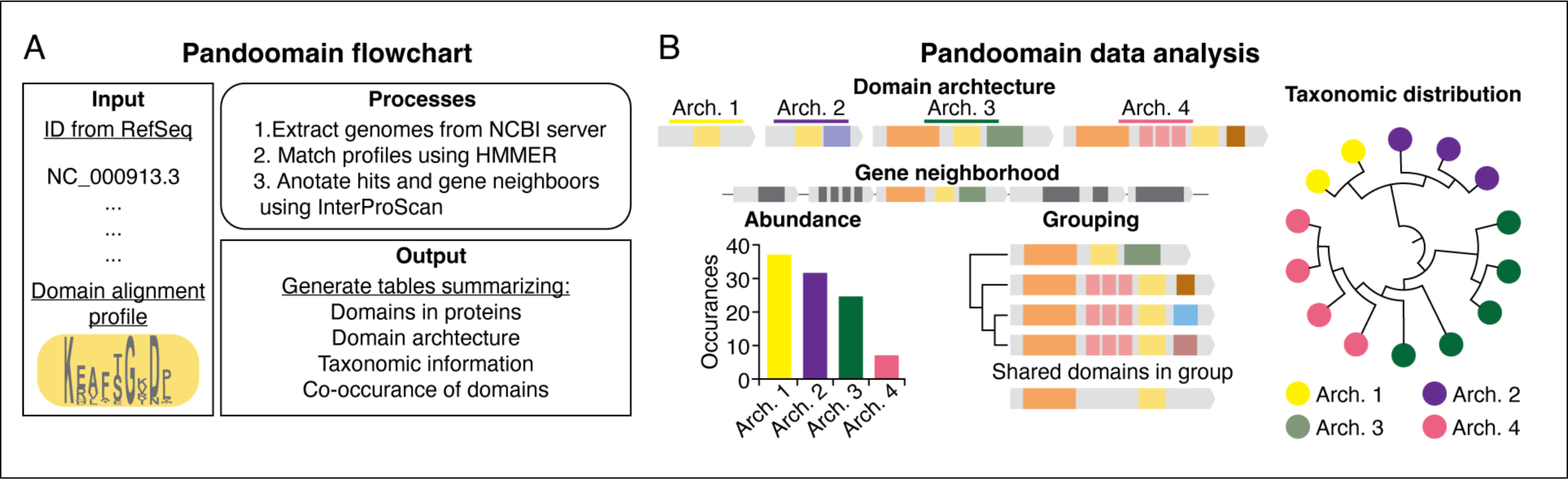
Overview of the Pandoomain workflow and data analysis pipeline. (A) The flowchart illustrates the main operational steps of Pandoomain. (B) Visualization of the output depicting domain architecture, gene neighborhood, the relative abundance of each architecture, the grouping of similar architectures based on shared domains, and their taxonomic distribution.

## Methods

### Pandoomain implementation and workflow

The Pandoomain workflow is managed by a Snakemake[21] pipeline that executes a series of modular scripts, with key parameters specified in a config.yaml file. The pipeline is initiated with a user-supplied list of NCBI RefSeq[22] genome assembly accessions. Using this list, Pandoomain automates the download of the corresponding protein FASTA and annotation (GFF) files for each genome from the NCBI Assembly database, storing them locally for subsequent analysis. Pandoomain downloads this pre-compiled database and merges it with the genome metadata, providing a file (genomes_ranks.tsv) with standardized and complete taxonomic information for the ranks superkingdom, phylum, class, order, family, genus, and species.

Once the genomic data is acquired, Pandoomain uses the HMMER homology search algorithm[23] to scan all downloaded proteomes against a user-provided directory of Hidden Markov Model (HMM) profiles. The HMM profiles can be generated by the user with the HMMER package, or downloaded from different databases. This step identifies all instances of the target protein families of interest, producing a detailed report (hmmer.tsv) of all hits, their genomic locations, and search scores.

Following the identification of target proteins, Pandoomain systematically analyzes the genomic neighborhood surrounding each hit. The size of this neighborhood is a user-configurable parameter, defaulting to 12 genes upstream and downstream. This analysis yields a catalog (neighbors.tsv) with every protein within these defined neighborhoods.

The pipeline then extracts the amino acid sequences of all identified proteins, both the initial HMMER hits and their neighbors, into a single multi-FASTA file (all.faa). This file is then processed in batches by InterProScan[13], a powerful tool for large-scale protein domain annotation.

The script processes the InterProScan output to define the domain architecture of each protein. Each identified domain is assigned a random character for downstream similarity analysis between domain architectures and to facilitate inspection by the user. Subsequently, the script synthesizes the taxonomic data, protein hit information, and domain annotations to generate a presence-absence matrix (absence_presence.tsv). This matrix provides a clear, high-level overview of the distribution of every identified protein domain across all input genomes, a format ideally suited for downstream phylogenetic profiling, co-evolutionary analysis, and statistical comparisons. The code require for Pandoomain implementation is available on https://github.com/deMoraes-Lab/Pandoomain.

### Defining domain architecture groups

To classify the different domain architectures into distinct groups, we used the single-character per domain representation of proteins to build a similarity matrix of all protein pairs using Jaccard index[24]. The R package ape[25] was then used to build a dendrogram of similarity using the UPGMA clustering method. The resulting dendrogram was then manually inspected using the iTOL web interface[26], and clades were grouped to represent the major architectural characteristics.

### Taxonomy analysis and visualization

To visualize the taxonomic distribution of the identified architecture groups, we used the Environment for Tree Exploration (ETE) Python toolkit[27]. Specifically, the ncbi_taxonomy module was used to build a phylogenetic tree based on the NCBI taxonomy IDs obtained from each genome’s metadata. The final tree, annotated with the domain architecture groups, was generated using the iTOL web interface.

### Identification of new W10XG toxins

To identify new W10XG toxins, proteins containing W10XG domains were filtered to remove sequences with fewer than 260 amino acids. This threshold was defined because it is in this approximate region that the PT-TG domain ends and the toxin domain starts. The putative toxin domains were extracted from the C-terminal portion of the proteins, from position 260 to the end, and aligned using MAFFT[28]. The toxins were then clustered into homology groups sharing more than 70% sequence identity. One random member of each group was selected for further analysis. Putative immunity proteins were identified from the genes immediately downstream of the toxin gene. The structure of the toxin and immunity multimer was modeled using the AlphaFold web server[29]. Resulting protein structures were individually curated, and toxin-immunity pairs where the toxin domain appeared truncated or there was no predicted interaction between toxin and immunity were excluded. Finally, the resulting toxin domains were aligned using the Mason tool available on Foldseek web server[30]. If structural homologues were identified, only one was selected for further analysis.

### Bacterial culture methods and plasmid construction

*E. coli* DH5α strains used in this study were cultivated in Lysogeny Broth (LB) at 37°C with shaking or on LB medium solidified with 1.5% agar. When necessary, antibiotics were supplied to the media in the following concentrations: gentamycin (15 μg ml−1) and trimethoprim (50 μg ml−1). Synthetic DNA fragments containing codon-optimized toxin and immunity protein genes were acquired from Twist Bioscience. The fragments were designed with terminal homology regions for cloning into expression vectors via Gibson assembly (NEBuilder HiFi DNA Assembly Master Mix). Toxin genes were cloned into the rhamnose-inducible vector pSCRhaB2[31], and cognate immunity genes were cloned into the IPTG-inducible vector pPSV39[32]. To prevent self-intoxication during the cloning process, toxin constructs were transformed into *E. coli* DH5α already carrying the plasmid for the corresponding immunity protein.

### Validation of W10XG toxins

To test the activity of the toxin-immunity pairs, overnight cultures of *E. coli* carrying the dual-plasmid expression system were diluted 1:100 into fresh LB medium. The cultures were grown with shaking at 37°C to an OD_600_ of approximately 0.6. Expression was then induced under three conditions: Toxin expression was induced with 3 mM rhamnose, co-expression of the toxin and immunity protein was induced with 3 mM rhamnose and 1 mM IPTG, and a third culture was left uninduced as a negative control. Three biological replicates were used. After a 90-minute incubation period, samples were collected, serially diluted, and plated onto LB agar. Plates were incubated overnight at 37°C, and the resulting colonies were counted to determine cell viability (CFU ml−1).

## Results

### Implementation

We opted to use a SnakeMake architecture due to its modularity, which ensures that all intermediate files, such as the HMMER results and neighbor lists, are also available for detailed inspection, offering researchers multiple avenues for hypothesis generation and follow-up analysis. SnakeMake also executes the required scripts without requiring user input, which streamlines the entire process. The RefSeq genome accessions can be easily extracted from the NCBI Dataset web page that allows the search of entries using the name of organisms or specific TaxIDs. The web page also allows filtering by assembly status, deposit year, and other parameters. Additionally, the NCBI Datasets command-line tools (CLI)[33] can be used to download large numbers of accessions. Pandoomain then automates the download of the protein FASTA and GFF annotation files for all the specified genomes from the NCBI Assembly database. After the conclusion of this step, all the data is locally available for subsequent, rapid processing. The Pandoomain analysis pipeline begins by identifying proteins that match a user-provided Hidden Markov Model (HMM) profile. This HMM alignment can be generated by the user from a custom set of amino acid sequences or acquired from public databases. The InterPro database is a valuable source for such profiles, as it integrates data from 13 different member databases, providing a comprehensive collection of protein signatures.

The integration of taxonomic and genomic data is a common challenge in functional genomic analysis. The NCBI taxonomic data, while comprehensive and well curated, is especially challenging for high-throughput analysis of genomes from diverse organisms. The data is organized in a tree topology where each taxon is assigned to up to 41 taxonomic ranks, and these ranks can be a gap in the lineage of many organisms, as standard ranks like ‘class’ or ‘order’ may be missing. By employing the Taxallnomy tool[34], we can obtain a hierarchically complete taxonomic table and obtain the complete taxonomic lineage with all ranks of taxa available in the NCBI Taxonomy database. As a result, we can determine the taxonomic relationship between all the organisms in the analysis.

A key feature of the pipeline is the automated analysis of the genomic neighborhood surrounding each protein identified by the HMM search. The pipeline defines the neighborhood as a user-configurable number of genes flanking the target gene, typically twelve upstream and twelve downstream. All proteins within this window are then annotated using HMMER against the Pfam database. This automated annotation of the genomic context leverages the principle of ‘guilt by association’ to make functional inferences.

### Pandoomain application to study protein containing PT-TG domain

Bacterial polymorphic toxins are composed of different trafficking domains associated with toxin domains. In many cases, they also contain pre-toxin domains[6]. This modular architecture results in a wealth of domain architectures distributed across all bacteria, different secretion systems, and employing a still not fully described arsenal of toxins.

We chose to use the pre-toxin TG domain (PT-TG, PFAM: PF14449) in our proof-of-concept application of Pandoomain because of its widespread distribution across bacterial antagonistic systems, its unresolved biological function, and its association with diverse domains[19].

The PT-TG domain named for its conserved TG motif, is hypothesized to function as a linker between toxin and trafficking domains or to mediate interactions with the target cell envelope[19]. To the best of our knowledge, the PT-TG domain itself remains unresolved, likely due to intrinsic disorder. To investigate its function, we generated an HMM alignment using the seed sequences from the PFAM database. This analysis showed that other residues, including a Gly, Asp, and Arg within a 15-amino-acid region, are more conserved than the nominal TG motif (Fig. 2A). We next used AlphaFold3 to model the structure of proteins containing a PT-TG domain. Consistent with its presumed disordered nature, this region was not predicted with high confidence (pLDDT < 70). Despite the overall low confidence, the models consistently predicted that the highly conserved 15-residue region forms a loop (Fig. 2B). The combination of high residue conservation and a consistent structural prediction of a loop suggest this region is a key functional element of the PT-TG domain, likely mediating molecular interactions required for toxin delivery.

**Fig 2.**
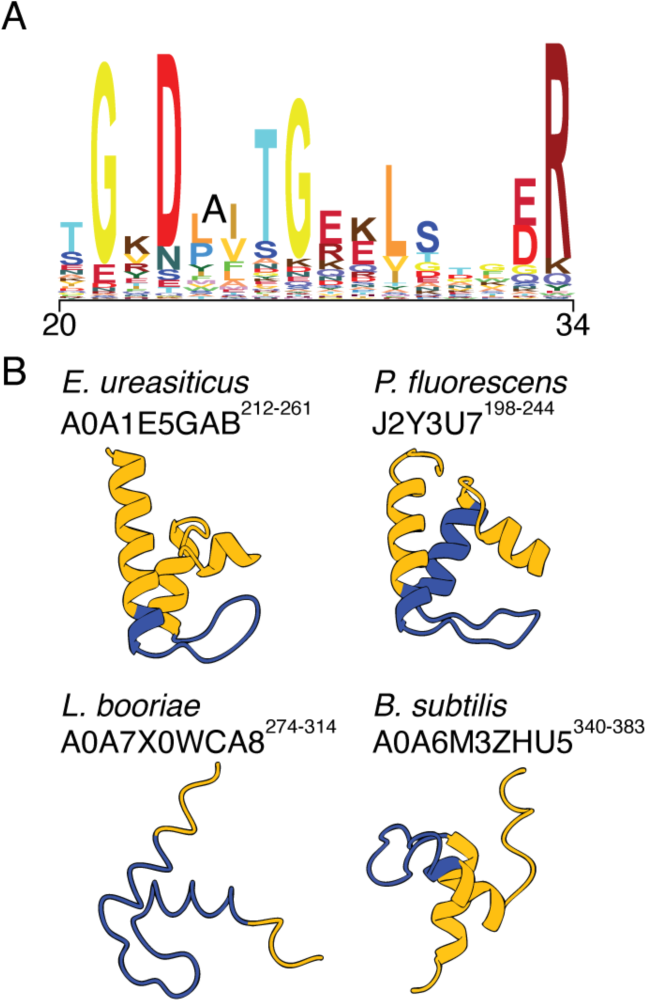
Conserved residues of PT-TG are in an unorganized region. (A) A sequence logo depicting the conserved 15-amino-acid region of the PT-TG domain shows that residues including a Gly, Asp, and Arg are more conserved than the nominal TG motif. (B) AlphaFold3 structural models of the PT-TG domain from four different bacterial proteins originating from *Enterococcus ureasiticus*, *Pseudomonas fluorescens*, *Listeria booriae*, and *Bacillus subtillis*. The models predict that the highly conserved 15-residue region lacks secondary structure.

### Classification of domain architectures containing PT-TG

Using an HMM query generated with the PT-TG alignment available on Interpro, we employed Pandoomain to search for homologs across ∼300,000 bacterial genomes (Table S1). The selected genomes were filtered based on the CheckM[35] completeness of >95% and contamination <0.5%. To reduce the presence of genomes from clonal strains, we selected only one genome per Bioproject deposition. This step reduces the amount of redundant data without the need to apply computationally intensive algorithms to identify nearly identical genomes over the >2.5M entries available on NCBI.

Using Pandoomain, we analyzed the selected genomes and identified 10,226 proteins containing the PT-TG domain distributed across 22,629 genomes, highlighting its widespread prevalence. These proteins showed remarkable architectural diversity, comprising 312 unique arrangements that combine the PT-TG domain with 111 other distinct protein domains (Table S2, Table S3).

The entire analysis was performed on a MacBook Pro (M1, 16 GB RAM), with a total runtime of approximately 19 hours. An external solid-state drive (SSD) was necessary to store the large number of genomic files downloaded. While all local computation steps completed without issue, the initial data acquisition was occasionally interrupted. Users should be aware that the download phase is dependent on the connection stability with NCBI servers, and interruptions can be a common issue when retrieving large datasets.

To manage the complexity of 312 unique domain architectures, we clustered them into a smaller number of functional groups using the Jaccard index. We chose this metric as it provides a robust measure of similarity based on the presence or absence of unique domains, which is particularly effective for this analysis since many toxins feature repeated domains within a single protein. This approach disregards the number of repeated domains within a given protein, allowing us to group architectures that share the same core trafficking domains, even if they are associated with different toxin domains.

This clustering approach successfully condensed the 312 architectures into 45 main groups (Fig. 3, Table S2). Our analysis revealed that the PT-TG domain is co-localized with tracking domains from a wide variety of known antagonistic systems, including the VgrG from the Type VI (T6SS)[36], WXG and LXG from the Type VII (T7SS)[37, 38], and filamentous hemagglutinin (FHA)-peptide repeat from contact-dependent inhibition (CDI) systems[39]. The architectures also incorporate diverse trafficking and repetitive domains, such as the choline-binding[40], cohesin[41], and dockerin[42] domains, which highlight their modular nature. A large fraction of the identified proteins did not contain any recognizable toxin domain or other domains previously associated with toxin delivery. This suggests that either these proteins utilize novel, uncharacterized toxin domains or they have evolved to serve functions distinct from direct interbacterial antagonism.

**Fig 3.**
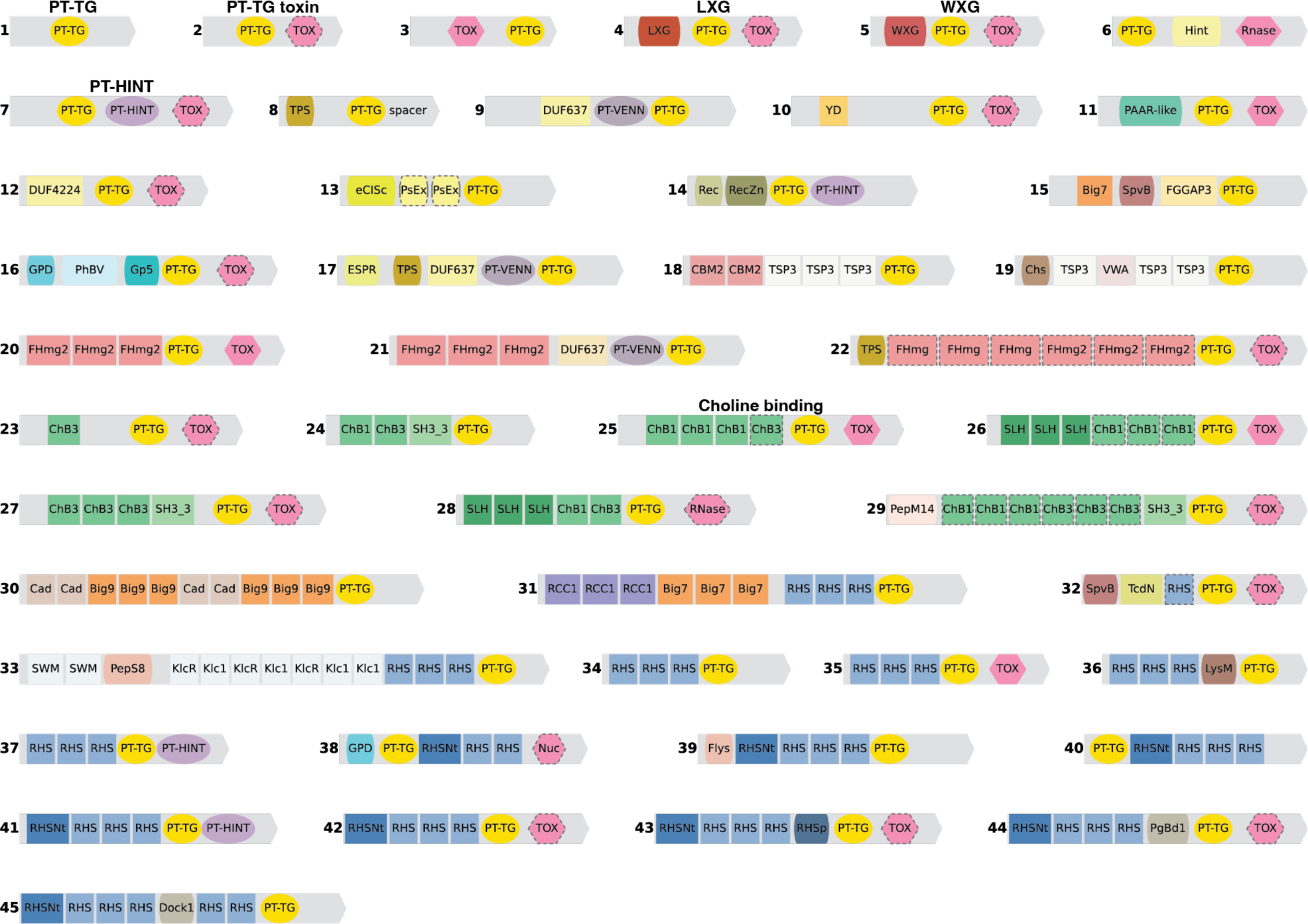
Pandoomain analysis reveals 45 major groups of pt-tg domain architectures. The 312 unique domain architectures containing a PT-TG domain were clustered into 45 main groups based on the Jaccard index similarity of their component domains. Each numbered bar represents the consensus architecture for a group, with different colored boxes corresponding to distinct protein domains. The architectures that include 95% of all proteins are named. The colored shapes indicate their function: Rectangle – repetitive domain, ellipse – pre-toxin, hexagon – toxin, and rounded rectangle – other. Dotted lines indicate domains that may be absent.

Notably, this analysis identified in architectures 30 and 31 the presence of Bacterial Ig-like (Big) domains 7 and 9 (PF17957 and PF17963) that were not previously associated with polymorphic toxin systems. The Big domains share a similar fold to immunoglobulins (Ig), are widespread in bacteria, and are involved in diverse roles that require interaction between different molecules[43]. Among their functions, these domains are in cell-surface proteins and contribute to the attachment to eukaryotic host cells, promoting pathogenesis[44]; they facilitate the attachment hydrolases to polysaccharide substrates[45, 46]; they may facilitate mating pair formation in conjugation[47]; and are present in phages and hypothesized to contribute to adsorption[48]. In architectures 30 and 31, Big domains appear in tandem repeats, suggesting they serve a structural role similar to that of RHS[49], ALF[50], and FHA[39] domains. Given that Big domains are frequently involved in cell-to-cell recognition, these domains may mediate the molecular interactions between donor and recipient cells during toxin translocation.

### Abundance and distribution of architecture groups

After grouping the architectures, we investigated their abundance and taxonomic distribution. The analysis revealed a highly skewed distribution: most architectures are rare, with 14 groups found to be singletons, containing one member, while a few are highly abundant (Fig. 4A). In contrast, just six architecture groups, LXG, PT-TG, PT-TG toxin, WXG, PT-HINT, and Choline-binding, collectively accounted for over 95% of all identified proteins containing a PT-TG domain. Among these, the LXG and WXG architectures are known components of the T7SS[37, 38]. The PT-HINT group contains a pre-toxin hedgehog-intein peptidase domain predicted to facilitate toxin release at the cell surface[19]. This group was found in many clades, most of its members were from the Streptomycetaceae family (Fig. 4B). We also noted that members of this architecture possess an unannotated N-terminal region of approximately 300 amino acids, suggesting they may utilize additional mechanisms for secretion and translocation. Taxonomic analysis showed that proteins with a PT-TG domain are most prevalent in the family Bacillaceae (Fig. 4B). The T7SS-associated LXG and WXG domains were found almost exclusively in monoderms. The few exceptions in Gammaproteobacteria and Alphaproteobacteria were located on small contigs lacking other genes and are likely assembly artifacts. The Choline-binding group was restricted to Bacillaceae, indicating it may be part of an antagonism system specific to this family. In stark contrast, the PT-TG singleton and PT-TG toxin groups were widely distributed across diverse bacterial phyla, including both monoderm and diderm lineages (Fig. 4B). Since antagonistic systems require specialized machinery to translocate toxins across distinct cell envelopes, this broad distribution suggests that these architecture groups are not a single, conserved system but associated with multiple, distinct secretion systems. The unexpected discovery of the PT-TG domain association with both monoderm and diderm bacteria opens new avenues for investigating the evolution and mechanisms of interbacterial competition.

**Fig 4.**
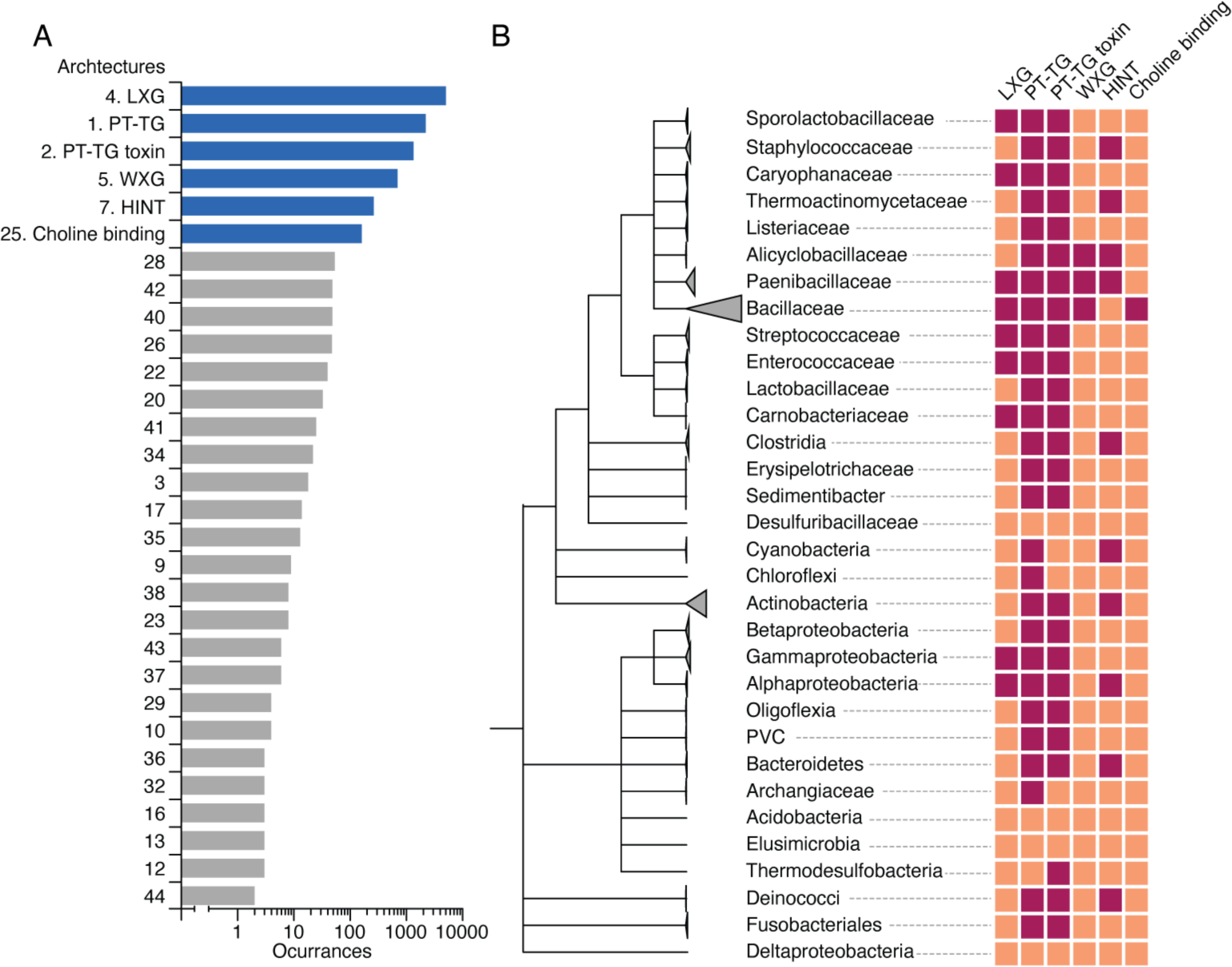
Abundance and taxonomic distribution of PT-TG domain architectures. (A) A bar chart showing the abundance of the PT-TG domain architecture groups with more than one member identified by Pandoomain. The top six most abundant groups, accounting for over 95% of all identified proteins, are highlighted in blue. (B) (B) A heatmap illustrates the taxonomic distribution of the six most abundant architecture groups across a phylogenetic tree of bacterial taxa. To account for the sparse representation of certain groups, branches of the taxonomic tree are collapsed at different hierarchical levels (e.g., family, class, or phylum).

### Genomic context analysis reveals a novel WXG domain variant

Our analysis revealed a large number of proteins containing a PT-TG domain that lacked any recognizable trafficking domain. This was an unexpected finding, which lead us to hypothesize that these proteins could represent truncated gene fragments. We compared the protein length distributions of the PT-TG singleton and PT-TG toxin groups against those of the well-characterized LXG and WXG groups. Proteins in the PT-TG singleton and PT-TG toxin groups have a wider length distribution than those in the LXG and WXG groups (Fig. 5A). Although some proteins are considerably smaller, particularly in the PT-TG singleton group, many members of both PT-TG architectures are over 250 amino acids long. While these observations do not exclude the possibility that some of these are truncated proteins, the substantial size of many members indicates that truncation alone cannot account for the general absence of a known trafficking domain or toxin domain.

**Fig 5.**
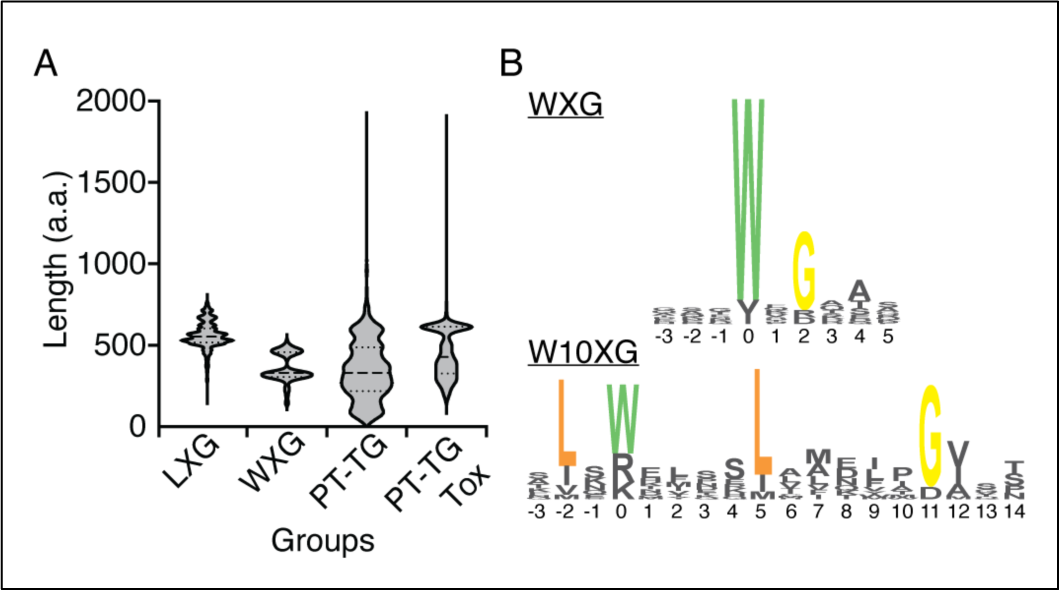
Characterization of the W10XG domain variant and its associated novel toxin families. (A) Violin plots comparing the amino acid (a.a.) length distributions of proteins from the LXG, WXG, PT-TG singleton, and PT-TG toxin architecture groups. The substantial size of many proteins in the PT-TG groups indicates that the absence of a known trafficking domain cannot be explained by truncation alone. (B) Sequence logo alignment comparing the canonical WXG domain with the W10XG variant identified in this study. The W10XG variant is defined by a characteristic 10-residue insertion between the conserved tryptophan (W) and glycine (G) residues.

To investigate why many PT-TG proteins lacked trafficking domains, we analyzed their genomic neighborhoods for functional clues. Using the Apriori algorithm[51] for association rule mining, we found that proteins from the PT-TG singleton and PT-TG toxin groups frequently co-occur with T7SS structural genes, including those encoding EsaE and WXG domains (PF16887 and PF06013). This led us to hypothesize that these proteins were not truncated but instead contained divergent LXG or WXG domains with homology below the detection threshold of InterproScan.

We performed a targeted HMMER search against the PT-TG singleton and PT-TG toxin proteins found in the genomic neighborhood of T7SS structural genes. This approach identified WXG domain homology in 112 proteins from the PT-TG singleton group and 95 from the PT-TG group. The bit scores for these hits were high but fell just below the Gathering Threshold (GA) defined by PFAM, explaining their initial omission. To understand the reason for the lower score, we aligned the N-terminal regions of these newly identified proteins with canonical WXG domains. The alignment revealed a distinct variation: a 10-residue insertion containing a conserved leucine between the characteristic tryptophan (W) and glycine (G) residues. An additional conserved leucine was also present at the −2 position relative to the tryptophan (Fig. 5B). While this variation does not warrant the creation of a new domain classification, for simplicity, we will refer to this group as W10XG throughout the remainder of this manuscript. The discovery of the W10XG variant underscores the potential for false negatives in automated annotation pipelines. It also highlights the importance of manual curation and an understanding of the specific domains being studied to correctly interpret large datasets.

### Novel toxins discovery and validation

We hypothesized that the absence of identifiable toxin domains in many W10XG proteins is due to the presence of novel toxins that lack significant homology to established protein families. To identify these putative toxins, we implemented a computational filtering pipeline. First, we excluded proteins shorter than 260 amino acids to remove potential truncations. Next, we used FoldSeek to search for structural homologs of known toxins. Finally, we refined this candidate list by selecting for proteins encoded immediately upstream of a putative immunity gene, a common genomic organization for toxin-immunity pairs. This criterion was further supported by structural modeling, which predicted a stable heterodimer interaction between the candidate toxin and its cognate immunity protein. This pipeline identified 73 putative toxins, which we subsequently clustered into 24 groups based on high sequence and structural homology (Fig. 6A).

**Fig 6.**
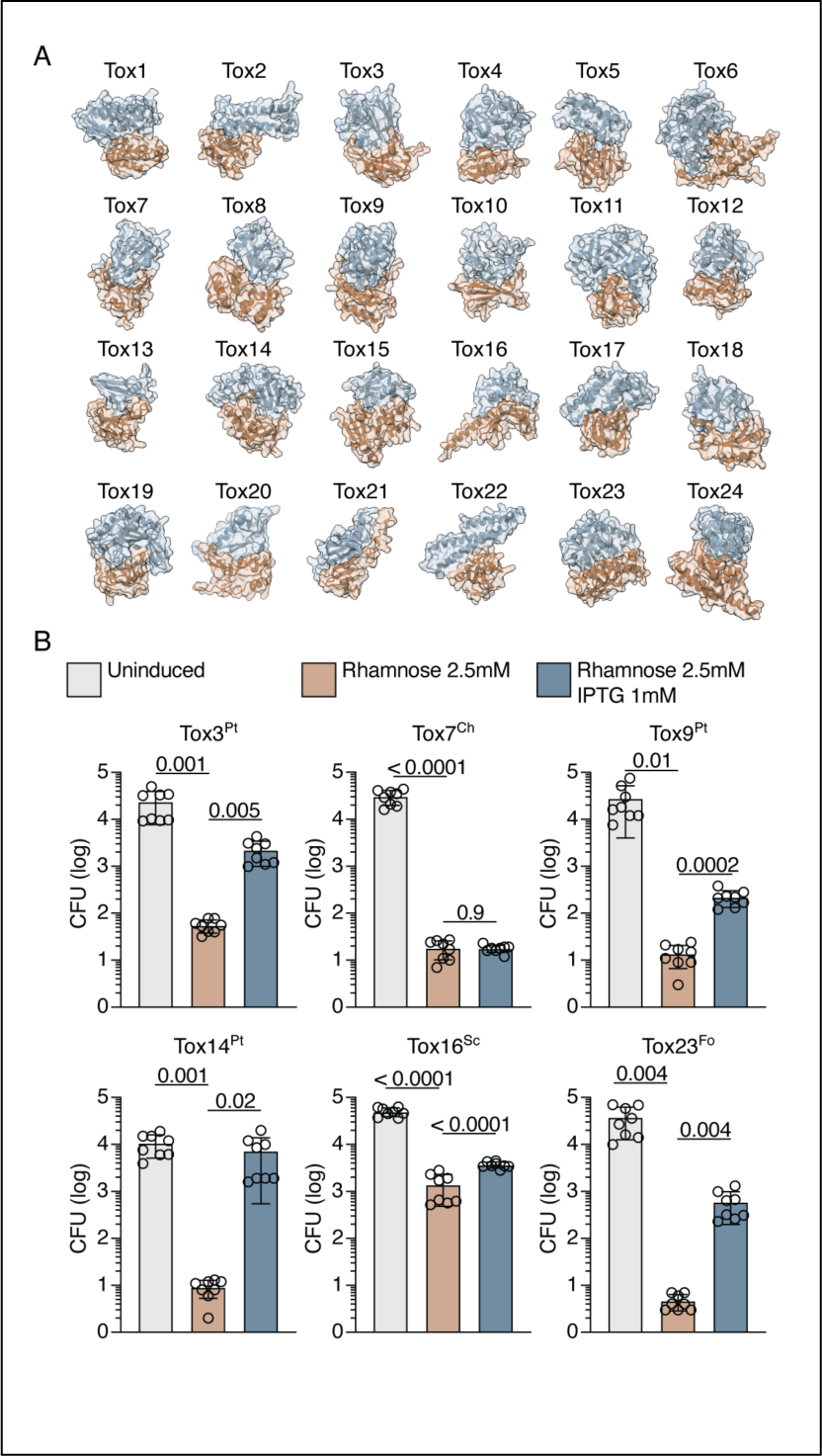
Structural diversity and experimental validation of novel W10XG-associated toxins. (A) AlphaFold-predicted structural models of a representative toxin-immunity complex from each of the 24 novel toxin groups (Tox1–Tox24) discovered in association with the W10XG domain. The putative toxin is shown in brown and the cognate immunity protein in blue. (B) Experimental validation of six representative toxin-immunity pairs. The bar charts show the viability of *E. coli* cells carrying a dual-inducible expression plasmid. Each dot represents a biological replicate. Cultures were either left uninduced (control), induced for toxin expression (2.5 mM rhamnose), or induced for co-expression of the toxin and its immunity protein (2.5 mM rhamnose and 1 mM IPTG). Statistical analysis was performed with one-way ANOVA. The p-value is indicated above the bars.

To experimentally validate these predictions, we selected six representative toxin-immunity pairs for functional analysis. We synthesized codon-optimized versions of each toxin and immunity gene and cloned them into a dual-inducible *E. coli* expression system, with toxin expression controlled by rhamnose and immunity protein expression by IPTG. Upon induction, all six putative toxins caused a significant reduction in cell viability, ranging from one to three orders of magnitude. Co-expression of the cognate immunity protein reduced this toxic effect in five of the six pairs, resulting in a significant recovery of viable cells. The one exception was the pair from Group 4, sourced from *Caldifermentibacillus hisashii*, for which the immunity protein failed to provide protection. This is potentially an artifact of the heterologous system, where issues such as low expression, improper protein folding, or degradation can occur[52]. These results provide strong experimental evidence that our pipeline successfully identified multiple groups of bona fide bacterial toxins.

### Mobile elements are enriched in the gene neighborhood of WXG and the variant W10G proteins

While investigating the gene neighborhood of W10XG proteins, we observed the presence of insertion sequences (IS) and other mobile elements in their gene neighborhood. To determine whether mobile elements are enriched in the gene neighborhood of W10XG proteins, we quantified the number of genomes where mobile elements were within 6 kilobases of different PT-TG groups. Specifically, we compared the association of the groups W10XG, WXG, and LXG with mobile elements due to their known biological function and total number. We found that W10XG and WXG proteins have approximately 40% of their members near mobile elements, in contrast to a statistically significant four-fold lower frequency for LXG proteins (Fig. 7A). The mobile elements associated with W10XG and WXG were diverse and belonged to different groups, including transposases from the families IS110, IS4, IS66, and TC1 to name a few. An in-depth search of PFAMs within the mobile elements revealed the presence of thirty-three different domains in the mobile elements, reinforcing their highly heterogeneous nature and suggesting that this is a region of diversification (Supplemental).

**Fig 7.**
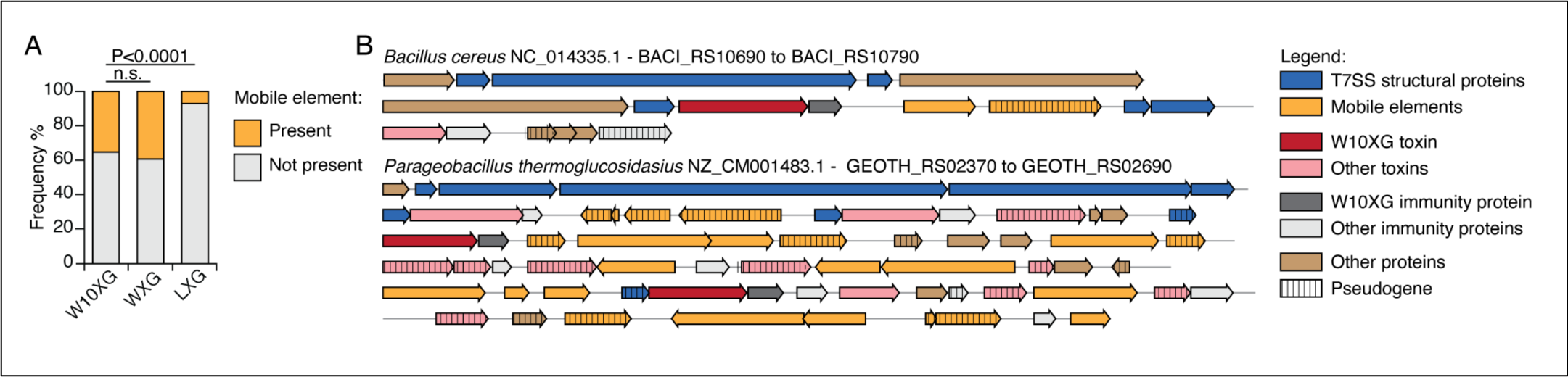
W10XG and WXG toxins are enriched in the vicinity of mobile genetic elements. (A) Stacked bar chart showing the frequency (%) of W10XG, WXG, and LXG proteins that have a mobile genetic element within a 6 kb genomic window. Both W10XG and WXG are significantly enriched in mobile elements compared to LXG. The P-value shown was calculated using a two-sided Fisher’s exact test; n.s., not significant. (B) Representative genomic loci from *Bacillus cereus* and *Parageobacillus thermoglucosidasius* illustrating the co-localization of W10XG toxin-immunity pairs with T7SS structural proteins, diverse mobile elements, and pseudogenes. Genes are colored according to the provided legend.

To exemplify the co-occurrence of WXG and W10XG proteins and mobile elements, we selected genomic regions of *Bacillus cereus* and *Parageobacillus glucosidasius* (Fig. 7B). In both cases, the regions contained genes encoding for T7SS structural proteins, toxin-immunity pairs, the mobile elements, and genes not related to T7SS or mobile elements. Also, the presence of pseudogenes was a common feature in these regions, which can be a result of active mobile elements resulting in gene decay [53]. Despite the similarities, the regions in *B. cereus* and *P. glucosidasius* also contained contrasting features, with the latter exhibiting a considerably larger number of toxins and mobile elements. Additionally, the W10XG gene neighborhood of *P. glucosidasius* contained mobile elements from different families, and several pseudogenes that appear to have been created as a result of truncations (Fig. 7B). ISs were the dominant mobile genetic element (MGE) found in the genomic region of PT-TG toxins. These mobile elements are composed exclusively of the DNA regions and enzymes required for self-mobilization[53]. Thus, it is unlikely that they are carrying PT-TG toxins across different genomes. One potential explanation for the observed enrichment of MGEs is that while mobilizing in the bacterial genome, they may disrupt or reactivate T7SS toxins, resulting in diversity of the repertoire of functional toxins in a population. A similar phenomenon was observed in T6SS toxins in *Salmonella enterica*, where recombination in the RHS repetitive domains resulted in a swap of active toxins, granting a competitive advantage to a subgroup of cells[54].

## Discussion

In this work, we developed and validated Pandoomain, a pipeline for scalable, context-based functional genomics that systematically downloads bacterial genomes, identifies proteins with user-defined domains via HMMER, and annotates the target proteins and their genomic neighbors using InterProScan. The command-line interface of Pandoomain is essential for its scalability and customizability. This design may be unfamiliar to researchers not used to computational workflows. By orchestrating the pipeline with Snakemake, the pipeline benefits from its inherent fault tolerance. Its execution verifies the existence of the expected output file for each task, allowing a seamless restart after interruption. This feature lowers the technical expertise required to run Pandoomain as it minimizes the need for user troubleshooting and manual intervention. In addition, we expect that command-line genomic tools will become more accessible with the increasing popularity of AI-powered assistant applications for translating natural language queries from users into specific commands.

A key differential of Pandoomain compared to other genomic mining tools is that it starts with the download of data, requiring only NCBI accession identifier. In its current version, the data acquisition is optimized for retrieval from the NCBI Assembly database. While this design choice provides reliable access to one of the largest public genomic resources, it represents a trade-off, as the tool does not yet natively support integration with other databases.

Pandoomain was designed with the explicit goal of analyzing thousands of genomes simultaneously, a scale at which generating graphical outputs directly from the pipeline would be computationally overwhelming and impractical for user interpretation. Consequently, the only viable output strategy is to produce a series of structured, plain-text files, such as tab-separated value (TSV) and FASTA files. We mitigate the absence of built-in visualizations by ensuring these outputs are organized in a way that is easily parsed, processed, and plotted. For example, representative gene neighborhoods of interest can be extracted for plotting, as in Fig. 3, the character representation of protein domain architecture can be used directly for estimation of the Jaccard index.

Our analysis of the pre-toxin TG (PT-TG) domain serves as a powerful demonstration of Pandoomain’s utility for large-scale, context-driven discovery. By applying the pipeline to 300,000 bacterial genomes, we identified over 10,000 proteins containing the PT-TG domain, which were organized into 312 distinct domain architectures. This remarkable diversity underscores the highly modular nature of polymorphic toxin systems. Clustering these architectures revealed that the PT-TG domain is associated with a wide variety of known antagonistic systems, including Type VI, Type VII, CDI systems, as well as novel domains not previously linked to toxin delivery, such as Bacterial Ig-like (Big) domains.

The incorporation of standardized taxonomic data was essential for these insights, allowing us to infer the distribution of these architectures across the bacterial tree of life. This revealed clear phylogenetic patterns, such as the restriction of the Choline-binding group to the Bacillaceae family and the broad distribution of PT-TG singleton groups across diverse monoderm and diderm lineages.

The investigation also highlights how Pandoomain can uncover novel biology that is missed by standard annotation methods. A large fraction of the identified PT-TG proteins initially appeared to lack any recognizable trafficking domain. By leveraging genomic context analysis, we found that these proteins frequently co-occurred with T7SS structural genes. This observation prompted a more targeted HMMER search, which successfully identified the divergent W10XG domain variant.

The capacity of Pandoomain for large-scale analysis is crucial for the discovery of low-frequency toxins like the ones associated with W10XG, which would likely remain undetected in smaller-scale searches. This discovery underscores the potential for false negatives in automated annotation pipelines and validates the power of using genomic context to guide more sensitive searches. Ultimately, Pandoomain implementation and our comprehensive study of the PT-TG domain offer valuable resources for the scientific community that can guide future research endeavors into microbial functional genomics and polymorphic toxin function and evolution.

## Acknowledgements

This research was supported by a research grant from the National Science Foundation (Award Number:2443063). We thank Dr. Kassidy Hebert for thoughtful comments on the manuscript.

